# Sleep disruption precedes forebrain synaptic Tau burden and contributes to cognitive decline in a sex-dependent manner in the P301S Tau transgenic mouse model

**DOI:** 10.1101/2023.06.07.544101

**Authors:** Shenee C. Martin, Kathryn K. Joyce, Kathryn M. Harper, Viktoriya D. Nikolova, Todd J. Cohen, Sheryl S. Moy, Graham H. Diering

## Abstract

**Background:** Sleep is an essential process that supports brain health and cognitive function in part through the modification of neuronal synapses. Sleep disruption, and impaired synaptic processes, are common features in neurodegenerative diseases, including Alzheimer’s disease (AD). However, the casual role of sleep disruption in disease progression is not clear. Neurofibrillary tangles, made from hyperphosphorylated and aggregated Tau protein, form one of the major hallmark pathologies seen in AD and contribute to cognitive decline, synapse loss and neuronal death.

Tau has been shown to aggregate in synapses which may impair restorative synapse processes occurring during sleep. However, it remains unclear how sleep disruption and synaptic Tau pathology interact to drive cognitive decline. It is also unclear whether the sexes show differential vulnerability to the effects of sleep loss in the context of neurodegeneration.

**Methods:** We used a piezoelectric home-cage monitoring system to measure sleep behavior in 3-11month-old transgenic hTau P301S Tauopathy model mice (PS19) and littermate controls of both sexes. Subcellular fractionation and Western blot was used to examine Tau pathology in mouse forebrain synapse fractions. To examine the role of sleep disruption in disease progression, mice were exposed to acute or chronic sleep disruption. The Morris water maze test was used to measure spatial learning and memory performance.

**Results:** PS19 mice exhibited a selective loss of sleep during the dark phase, referred to as hyperarousal, as an early symptom with an onset of 3months in females and 6months in males. At 6months, forebrain synaptic Tau burden did not correlate with sleep measures and was not affected by acute or chronic sleep disruption. Chronic sleep disruption accelerated the onset of decline of hippocampal spatial memory in PS19 males, but not females.

**Conclusions:** Dark phase hyperarousal is an early symptom in PS19 mice that precedes robust Tau aggregation. We find no evidence that sleep disruption is a direct driver of Tau pathology in the forebrain synapse. However, sleep disruption synergized with Tau pathology to accelerate the onset of cognitive decline in males. Despite the finding that hyperarousal appears earlier in females, female cognition was resilient to the effects of sleep disruption.

## Background

Sleep is an essential conserved behavior that supports higher cognition such as learning and memory^1^. Declines in sleep amount and quality are expected components of aging and may contribute to the susceptibility of individuals to develop neurodegenerative diseases such as Alzheimer’s disease (AD)^2^. Once AD progression is underway, sleep continues to worsen, and sleep disruption tracks with the progression of cognitive decline and accumulation of neuropathology^3^, suggesting that sleep disruption may play a causal role in disease progression. An emerging body of work is suggestive of a bidirectional relationship linking sleep disruption and AD-pathology, where sleep disruption promotes pathology, and worsening pathology drives sleep disruption in a vicious feed-forward cycle^4–6^. Promoting restorative sleep may therefore be an important therapeutic strategy to halt or delay pathology progression and protect cognitive function. Neuronal synapses in the forebrain undergo widespread and profound remodeling during the sleep/wake cycle, or in response to sleep disruption, suggesting that synapses are a locus for the restorative benefits of sleep and a site of dysfunction in response to sleep loss^7–11^. Synapse dysfunction is well documented to occur in the progression of AD and is strongly linked with the emergence of cognitive decline^12^. Understanding the causes and consequences of sleep disruption, the links between sleep disruption and synapse dysfunction, and the time course of sleep disruption in relation to other aspects of AD progression will all be essential in developing the most effective sleep-based treatments for use in AD.

Age-dependent accumulation of extracellular amyloid plaques (Aβ) and intracellular neurofibrillary tangles (NFTs) made up of aggregate prone hyperphosphorylated Tau are hallmarks of AD. Tau pathology is also prevalent in several other neurodegenerative conditions such as frontal temporal dementia (FTD). Tau is a microtubule binding protein primarily localized to the neuronal axon where it supports axonal microtubule-based transport and structure^13–15^. During disease progression, phosphorylated Tau has been shown to re-localize to dendrites and post synaptic compartments, where it contributes to synapse dysfunction and synapse loss^16^. Synaptic Tau has been suggested to contribute to cognitive decline in the early stages of AD, in advance of robust neuron loss^16^. However, the function of Tau at synapses, and the links with sleep disruption are not fully understood. Pathological Tau-species are known to propagate between neurons and throughout the brain through synaptic connections and the extracellular space^17–20^. Tau spread, and propagation has been linked to sleep, where sleep disruption promotes Tau aggregation and spread in a neuronal activity dependent manner, and restorative sleep serves to promote Tau clearance^21^. Taken together, these results make a persuasive argument to focus on understanding the pathological role of synaptic Tau in AD and sleep disruption.

Recent studies have examined sleep behavior and the consequences of experimentally induced sleep disruption in a mouse model of AD/FTD that expresses an aggregation prone disease causing variant of human Tau, hTau-P301S. P301S mice, also called PS19, express disease linked Tau broadly across the brain from a transgene driven by the neuron specific PrnP (prion protein) promoter^22^. AD/FTD PS19 model mice have been shown to exhibit age-related Tau aggregation, neuroinflammation, synapse dysfunction, cognitive decline, and eventual neuron loss and brain atrophy^22, 23^. Age-related disease progression in these mice can be broadly described as pre-symptomatic at 3months of age, early-stage disease at 6months (sparse Tau pathology and minor changes in behavior), symptomatic disease stage at 9months (widespread pathology and robust cognitive decline but without widespread neurodegeneration), and end-stage disease at 11-12months (robust neuron loss, brain atrophy, and death)^22, 23^. Declines in rapid-eye movement (REM) sleep and non-REM (NREM) sleep, and reduced sleep bout duration (sleep fragmentation) has been reported in male PS19 mice in the symptomatic disease phase beginning at 9-11 months of age^24^. Sleep amount was found to negatively correlate with Tau pathology in sleep promoting areas of the brainstem, the sublaterodorsal area and parafacial zone, but not in the cortex, suggesting that Tau burden in sleep promoting brain areas was a driver of age-related sleep disruption^24^. Tau burden has also been observed in younger PS19 mice in the wake promoting region of the brain stem, the noradrenergic locus coeruleus (LC)^21, 25^. Indeed, LC is one of the first brain regions to show Tau pathology in humans and animal models, well in advance of many disease symptoms^26, 27^. Chronic experimentally induced sleep disruption in PS19 mice drove increased Tau burden in the LC, and subsequently accelerated the loss of LC noradrenergic neurons suggesting that sleep disruption is an important driver of pathology^25^. These studies highlight the important interaction between sleep disruption and pathology in the context of AD, but the question remains, does sleep disruption drive pathology, or does pathology drive sleep disruption? Biological sex is also a critically important factor to understand the role of sleep disruption in the progression of AD and Tau pathology. Males and females have been shown to have differential likelihood of developing neurodegenerative conditions^28^, and also have been found to respond differentially to sleep deprivation^29^, suggesting that there may be sex-specific vulnerabilities to the negative consequences of sleep loss.

Building from these recent studies, we wished to further characterize age-related changes in sleep behavior in PS19 mice of both sexes, and to test whether a causal relationship exists between sleep disruption and Tau burden in forebrain synapses, structures believed to be a major site of sleep function supporting cognition^1^. We also further tested the role of sleep disruption as a driver of cognitive decline in PS19 mice of both sexes. We report that PS19 mice of both sexes exhibit age-related loss of sleep during the dark phase, a “hyperarousal phenotype”, with an earlier onset in females, that precedes robust Tau pathology. As PS19 mice of both sexes age into the later symptomatic phases of disease we observe fragmentation of sleep, consistent with a prior report^24^. Thus, changes in sleep behavior occur as an early symptom in advance of robust pathology, but sleep quality continues to worsen with disease progression. We find that sleep behavior was not correlated with Tau burden in forebrain synapses, and forebrain synaptic Tau burden was not increased following acute or chronic sleep disruption treatments in either sex, suggesting that sleep disruption is not a direct driver of forebrain Tau pathology in the synapse. These results lead us to suggest that additional factors may form intermediate links between sleep disruption and synaptic Tau burden. Finally, chronic sleep disruption accelerated decline of hippocampal spatial memory in male, but not female PS19 mice. We conclude that sleep disruption is an early symptom of disease progression in this Tau-based disease model, and that sleep disruption plays a causal role in cognitive decline in a sex-dependent manner.

## Methods

### Mice

Animal procedures were all approved by Institutional Animal Care and Use Committee of the University of North Carolina (UNC) and performed according to guidelines set by the U.S. National Institutes of Health. P301S (PS19) male mice were obtained from the Cohen laboratory at UNC and bred with C57Bl/6J females purchased from Jackson Labs. C57Bl/6J females were allowed to acclimate to the housing at UNC for at least 2 weeks before breeding. Experimental animals were bred by crossing wild-type (WT) C57Bl/6J females with P301S positive males and housed in the animal facility until sleep behavioral or molecular analysis. Experiments were performed using 3-, 6-, 9-, and 11-month-old mice. Experiments include WT and P301S littermates of both sexes. WT breeders for our colony were replaced every 3 months with mice supplied from Jackson Labs.

### Sleep phenotyping and behavior analysis

PS19 and C57Bl/6J littermates were moved to our wake/sleep behavior satellite facility on a 12 h:12 h light:dark cycle (lights on 7 am to 7 pm). Individual mice were housed in 15.5 cm^2^ cages with bedding, food, and water. Before the beginning of data collection, mice were allowed to acclimate to the environment for at least two full dark cycles. No other animals were housed in the room during these experiments. Sleep and wake behavior were recorded using a noninvasive home-cage monitoring system, PiezoSleep 2.0 (Signal Solutions, Lexington, KY). As previously described^30^. Briefly, the system uses a Piezoelectric polymer film to quantitatively assess sleep/wake cycles, total amount of sleep and quality from mechanical signals obtained from breath rate and movement. Specialized software (SleepStats, Signal Solutions, Lexington, KY) uses an algorithm to discern sleeping respiratory patterns from waking respiratory patterns. Sleep was characterized according to specific parameters in accordance with the typical respiration of a sleeping mouse. Additional parameters were set to identify wake including the absence of characteristic sleep signals and higher amplitude, irregular signals associated with volitional movements, and even subtle head movements during quiet wake. Data collected from the cage system were binned over specified time periods: 1 h bins to generate a daily sleep trace, 12 h bins for average light- or dark-phase percent sleep or sleep bout lengths. Sleep bout length was determined by a 30-s interval contains greater than 50% sleep and terminates when a 30-s interval has less than 50% sleep. This algorithm has been validated in adult mice by using electroencephalography, electromyography, and visual evaluation^31–34^ and utilized in additional studies^21, 35^.

### Post synaptic density preparation

Male and female 3-, 6-, 9-, and 11-month mice were sacrificed, mouse whole cortex or hippocampus, were dissected in ice-cold phosphate buffered saline, and then frozen on dry ice, and kept at −80°C until further processing. Frozen mouse cortices were homogenized using 12 strokes from a glass homogenizer in ice-cold homogenization solution (320 mM sucrose, 10 mM HEPES, pH 7.4, 1 mM 2,2’,2’’,2’’’-(Ethane-1,2-diyldinitrilo) tetraacetic acid [EDTA], 5 mM Na pyrophosphate, 1 mM Na3 VO4, 200 nM okadaic acid [Roche]). Brain homogenate was then centrifuged at 1000 × g for 10 min at 4°C to obtain the P1 (nuclear) and S1 (postnuclear) fractions. The S1 fraction was then layered on top of a discontinuous sucrose density gradient (0.8, 1.0, or 1.2 M sucrose in 10 mM 2-[4-(2-hydroxyethyl)piperazin-1-yl]ethanesulfonic acid [HEPES], pH 7.4, 1 mM EDTA, 5 mM Na pyrophosphate, 1 mM Na3 VO4, 200 nM okadaic acid, 50 nM JZL195, protease inhibitor cocktail [Roche]) and then subjected to ultra-centrifugation at 82 500 × g for 2 h at 4°C. Material accumulated at the interface of 1.0 and 1.2 M sucrose (synaptosomes) was collected. Synaptosomes were diluted using 10 mM HEPES, pH 7.4 (containing protease, phosphatase, and lipase inhibitors) to restore the sucrose concentration back to 320 mM. The diluted synaptosomes were then pelleted by centrifugation at 100 000 × g for 30 min at 4°C. Synaptosomes were then resuspended in 50mM HEPES pH 7.4 then equal parts 1X triton (both containing inhibitors) was added. Samples were incubated for 10 minutes and subjected to ultra-centrifugation at 25 x g for 20 minutes at 4°C. The supernatant was then removed resulting in an enriched post synaptic fraction pellet. The pellet was then resuspended in 50mM HEPES, pH 7.4 (containing inhibitors).

Frozen mouse hippocampi were homogenized 15 times using a 28g needle in ice-cold homogenization solution mentioned above. Brain homogenate was then centrifuged at 1000 × g for 10 min at 4°C to obtain the P1 (nuclear) and S1 (postnuclear) fractions. The S1 fraction was then further centrifuged at 100 x g for 20 minutes at 4°C. The resulting S2 (cytosolic) fraction was aspirated leaving the P2 fraction which was resuspended in water (containing inhibitors) and incubated (via rotation) for 30-45 minutes at 4°C. Samples were then centrifuged at 25 x g for 20 minutes at 4°C. Crude synaptosomes were then resuspended in 50mM HEPES pH 7.4 then equal parts 1X triton (both containing inhibitors) was added. Samples were incubated for 10 minutes and subjected to ultra-centrifugation at 25 x g for 20 minutes at 4°C. The supernatant was then removed resulting in an enriched post synaptic fraction pellet. The pellet was then resuspended in 50mM HEPES, pH 7.4 (containing inhibitors).

The protein concentration was determined using Bradford assay and western blot samples were made up based on concentration averages to a known concentration.

### Western Blot

For Western Blot, proteins (5ug) were separated via sodium dodecyl sulfate polyacrylamide electrophoresis (SDS-PAGE) on a 10% acrylamide gel. After sample separation the proteins were transferred to 0.2-um nitrocellulose membrane (Bio-Rad) and incubated with 3% Bovine Serum Albumin (BSA; Fischer bioreagents) blocking buffer (100-mM Tris pH 7.5, 165-mM NaCl; TBS) for 30 minutes at room temperature. Membranes were then incubated with primary antibodies (full list provided in table 1) in a solution of 3% w/v BSA in TBST (100-mM Tris pH 7.5, 165-mM NaCl, 0.1% v/v Tween 20; TBST) at 4°C overnight. Membranes were then probed with secondary antibodies 1:15000 IRDye 680RD-conjugated goat anti-mouse (LICOR Biosciences) and 1:15000 IRDye 800RDconjugated donkey anti-rabbit (LI-COR) in 3% w/v BSA in TBST for 1 hour at room temperature. The immunoreactivity of all antibody signals was detected simultaneously with a Li-Cor Odyssey CLx IR imaging system (Li-Cor).

### Antibodies

**Table 1.** Western blot antibodies used in PS19 synaptic analysis

| Antibody | Species | Company | Concentration |
| --- | --- | --- | --- |
| Anti-AT8 | Mouse | Thermo Fisher | 1:1000 |
| Anti-AT180 | Mouse | Thermo Fisher | 1:1000 |
| Anti-AT100 | Mouse | Thermo Fisher | 1:1000 |
| Anti-pS396 | Rabbit | Thermo Fisher | 1:1000 |
| Anti-Tau1 | Mouse | EMD Millipore | 1:1000 |
| Anti-K9 | Rabbit | Agilent | 1:5000 |
| Anti-PSD95 | Mouse | NeuroMab 75-028 | 1:1000000 |
| Anti-GluA1 | Mouse | NeuroMab 75-327 | 1:1000 |
| Anti-pS845 | Rabbit | EMD Millipore | 1:1000 |
| Anti-pS831 | Rabbit | EMD Millipore | 1:1000 |
| Anti-GluA2 | Mouse | EMD Millipore | 1:1000 |
| Anti-GluA3 | Rabbit | Cell Signaling | 1:1000 |
| Anti-Arc | Mouse | Synaptic Systems | 1:1000 |
| Anti-GluN2A | Rabbit | Cell Signaling | 1:1000 |
| Anti-GluN2B | Rabbit | Cell Signaling | 1:1000 |
| Anti-GluN1 | Rabbit | Cell Signaling | 1:1000 |
| Anti-Hsc70 | Mouse | EMD Millipore | 1:1000 |

### Sleep disruption paradigms

For acute and chronic sleep disruption experiments, animals were brought up from the animal facilities either to our sleep recording satellite facility or behavior satellite room. Animals were acutely sleep deprived at 6 months via gentle handling for 4 hours as previously described^36^. Briefly, gentle handling consists of light tapping on home cages, rustling or removing bedding, replacing bedding, or movement on a cart. The mice remain in their home cage and are never touched. For chronic sleep disruption the animal’s home cage was placed on top of orbital shakers which automatically agitated at 110 rpm for 10 secs followed by still rest for 99 secs (109 secs total per cycle). This paradigm has been described by other groups noted to produce a mild fragmentation of sleep^25, 35, 37^. Control groups were housed alongside treatment groups atop inactive orbital shakers.

### Morris water maze behavioral paradigm

The water maze was used to assess spatial learning, swimming ability, and vision. The water maze consisted of a large circular pool (diameter = 122 cm) partially filled with water (45 cm deep, 24-26°C), located in a room with numerous visual cues. The procedure involved two phases: a visible platform test and acquisition in the hidden platform task.

Visible platform test. Each mouse was given 4 trials per day, across 2 days, to swim to an escape platform cued by a patterned cylinder extending above the surface of the water. For each trial, the mouse was placed in the pool at 1 of 4 possible locations (randomly ordered), and then given 60 sec to find the visible platform. If the mouse found the platform, the trial ended, and the animal was allowed to remain 10 sec on the platform before the next trial began. If the platform was not found, the mouse was placed on the platform for 10 sec, and then given the next trial. Measures were taken of latency to find the platform and swimming speed via an automated tracking system (Noldus Ethovision).

Acquisition of spatial learning in a hidden platform task. Following the visible platform task, mice were tested for their ability to find a submerged, hidden escape platform (diameter = 12 cm). Each mouse was given 4 trials per day, with 1 min per trial, to swim to the hidden platform. Criterion for learning was an average latency of 15 sec or less to locate the platform. Mice were tested until the group reached criterion, with a maximum of 9 days of testing. When the group reached criterion (on day 4 in the present study), mice were given a 1-min probe trial in the pool with the platform removed. Selective quadrant search was evaluated by measuring the percent of time spent in the quadrant where the platform (the target) had been located during training, versus the opposite quadrant, and by number of swim path crosses over the location (diameter = 12 cm) where the platform had been placed during training, versus the corresponding area in the opposite quadrant.

### Statistical Analysis

All data from sleep behavior experiments were analyzed in Microsoft Excel or GraphPad Prism version 9.1.0 (GraphPad Software LLC). To compare sleep amounts and bouts between PS19 and WT mice, unpaired student’s t-tests with Bonferroni correction for multiple comparisons were used. Behavioral tests were performed by experimenters blinded to mouse genotype. Statview (SAS, Cary, NC) was used for data analysis. One-way or repeated measures analysis of variance (ANOVA) were used to determine effects of genotype. Post-hoc comparisons were conducted using Fisher’s Protected Least Significant Difference (PLSD) tests only when a significant F value was found in the ANOVA. Within-genotype comparisons were conducted to determine quadrant selectivity in the water maze. All data is presented as mean ± SEM, *p<0.05, **p<0.01, ***p<0.001.

## Results

### Hyperarousal phenotypes emerge as an early symptom of disease progression in PS19 mice, particularly in females

Sleep disturbances are widespread in AD patients and have been reported in various animal models of AD, including male PS19 Tau transgenic AD/FTD model mice^4, 5, 24^. In order to better understand the causal links between Tau pathology and sleep disruption, and to ask whether there may be sex-specific vulnerabilities to sleep disruption, we first investigated the onset and progression of sleep disruption in PS19 mice by measuring sleep behavior at 3 months (pre-pathology), 6 months (early pathology), 9 months (late pathology, early atrophy) and 11 months (late pathology, late atrophy)^22, 23^. Sleep behavior was examined in male and female PS19 mice, in comparison to WT littermates, using the PiezoSleep Mouse Behavioral Tracking System (Signal Solutions), which allows for noninvasive monitoring of sleep/wake in a controlled light/dark environment. This system uses piezoelectric sensors to measure home-cage mouse motion and breathing to provide accurate measurements of total sleep time and bout lengths, achieving ∼95% accuracy when compared to EEG/EMG and has further been validated with video recordings^31–34^. Mice were maintained on a 12hr light/dark cycle and allowed a 2-day acclimation period to the home-cage recording system prior to data collection. Average daily total sleep amount and bout lengths were calculated from 13 consecutive days, providing robust measures of typical daily sleep patterns for each individual (Fig. 1A). As expected, all mice examined spent more time sleeping during the light phase and are more active during the dark phase. Overall, male and female WT and PS19 mice show comparable sleep amount in the light phase at 3- 6- 9- and 11 months, with the exception of PS19 females that showed a small but significant increase in light phase sleep amount at 3months (Fig. 1B,C; E,F). In contrast, sleep amount in the dark/active phase was significantly reduced in PS19 females as early as 3months and progressed with age through 11months (Fig. 1B,C). Even with the small increase in light phase sleep in females at 3months, the loss of dark phase sleep resulted in a significant decrease in total daily (24hr) sleep compared to WT controls at all ages (Figure 1C). Males showed a similar reduction in dark phase sleep that only became apparent at 6months of age and was progressive through 11months, again resulting in a significant reduction in total 24hr sleep (Fig. 1F). We refer to this selective loss of sleep during the dark phase as a “hyperarousal” phenotype. Sleep bout lengths, a crude measure of sleep quality, showed some significant reductions or compelling trends in males and females at 9-11 months, signifying reduced sleep quality with age and disease progression (Fig 1 D, G), consistent with a previous report from PS19 males^24^. Together, we find that dark phase hyperarousal, and consequent loss of total daily sleep, is an early symptom in PS19 mice and females particularly, and that sleep quality continues to decline with age.

**Figure 1.**
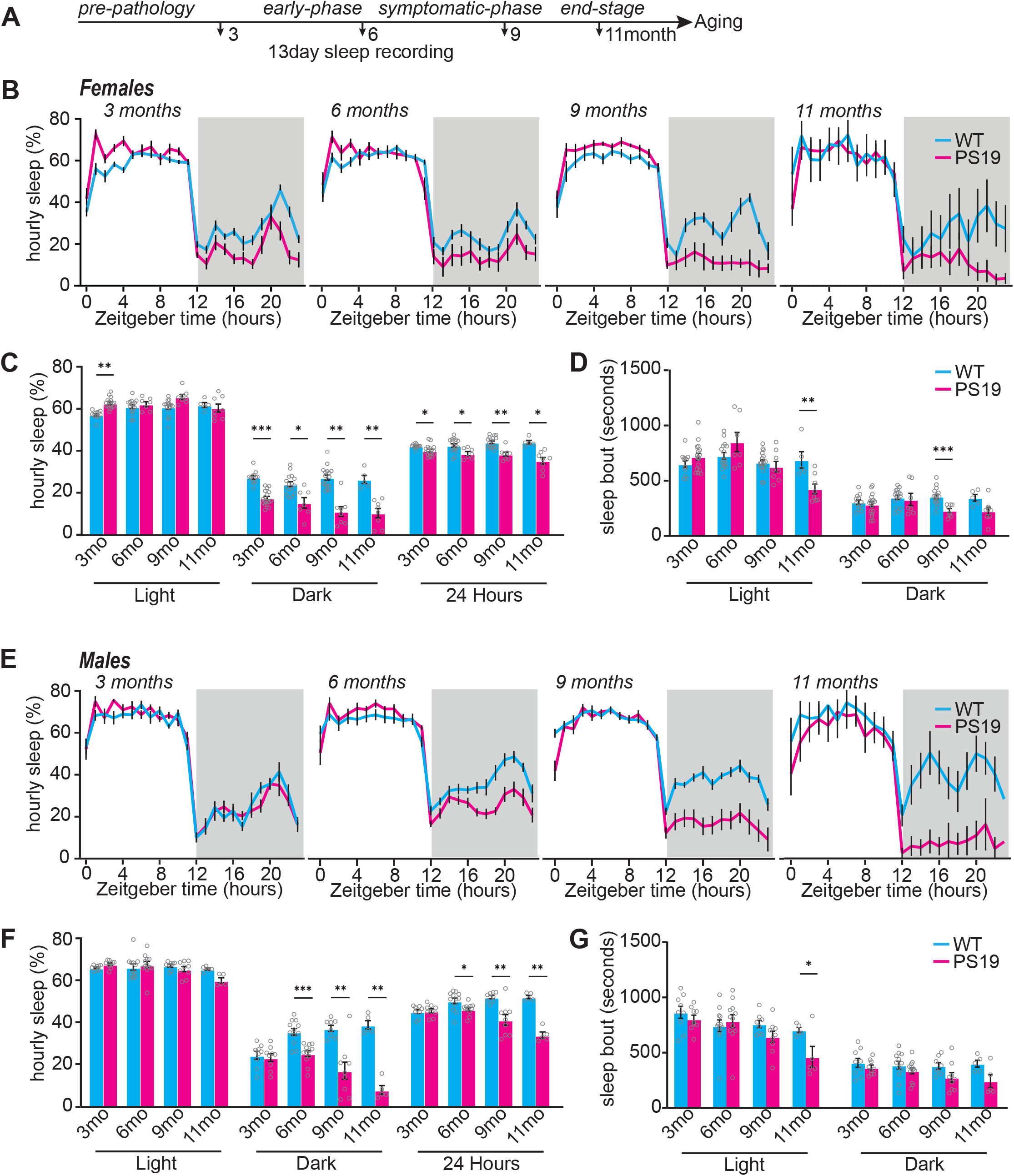
PS19 mice have decreased sleep that worsens with age. (A) experimental design (B) 24hr trace of average hourly sleep of female WT (blue line) PS19 (pink line) mice at 3 months (prepathology), 6 months (early-phase) 9 months (symptomatic-phase) and 11 months (end-stage). Grey bars in sleep traces indicate dark phase. (C and D) Quantification of average hourly sleep (C) and sleep bout length in seconds (D). Data separated into 12hrs of dark and light phases. (E) 24hr trace of average hourly sleep of male WT (blue line) and PS19 (pink line) mice at 3, 6, 9, and 11months. (F and G) Quantification of average hourly sleep (F) and sleep bout length in seconds (G). Data separated into 12hrs of dark and light phases. N=5- 15/age/sex/genotype. *p<0.05, **p<0.01, ***p<0.001 Unpaired two-tailed student’s t-test. Error bars indicate ± SEM.

### Tau burden in the forebrain synapse fraction begins to appear at 6months in PS19 mice

Forebrain synapses have been shown to be modified during sleep, mediating the benefits of sleep on cognitive function^1, 7^. Tau is normally an axonal protein but has been shown to be present in dendrites and the post synaptic density (PSD)^16^, where Tau contributes to synapse dysfunction and cognitive decline. The PSD is a fraction highly enriched with synaptic proteins and is amenable to biochemical isolation^7^. A recent publication has shown that phospho-Tau (AT8 epitope) becomes enriched in the hippocampal PSD fraction of 9-month-old PS19 females^23^. However, the link between forebrain synaptic Tau pathology and sleep disruption is unclear. We used biochemical fractionation and Western blot to assess the presence of pathological Tau (AT8 positive) in the PSD factions isolated from cortex of WT and PS19 mice at 3, 6, and 9months (Fig. 2A). No AT8-Tau was detected in WT littermates at any age (Fig. 2B).

**Figure 2.**
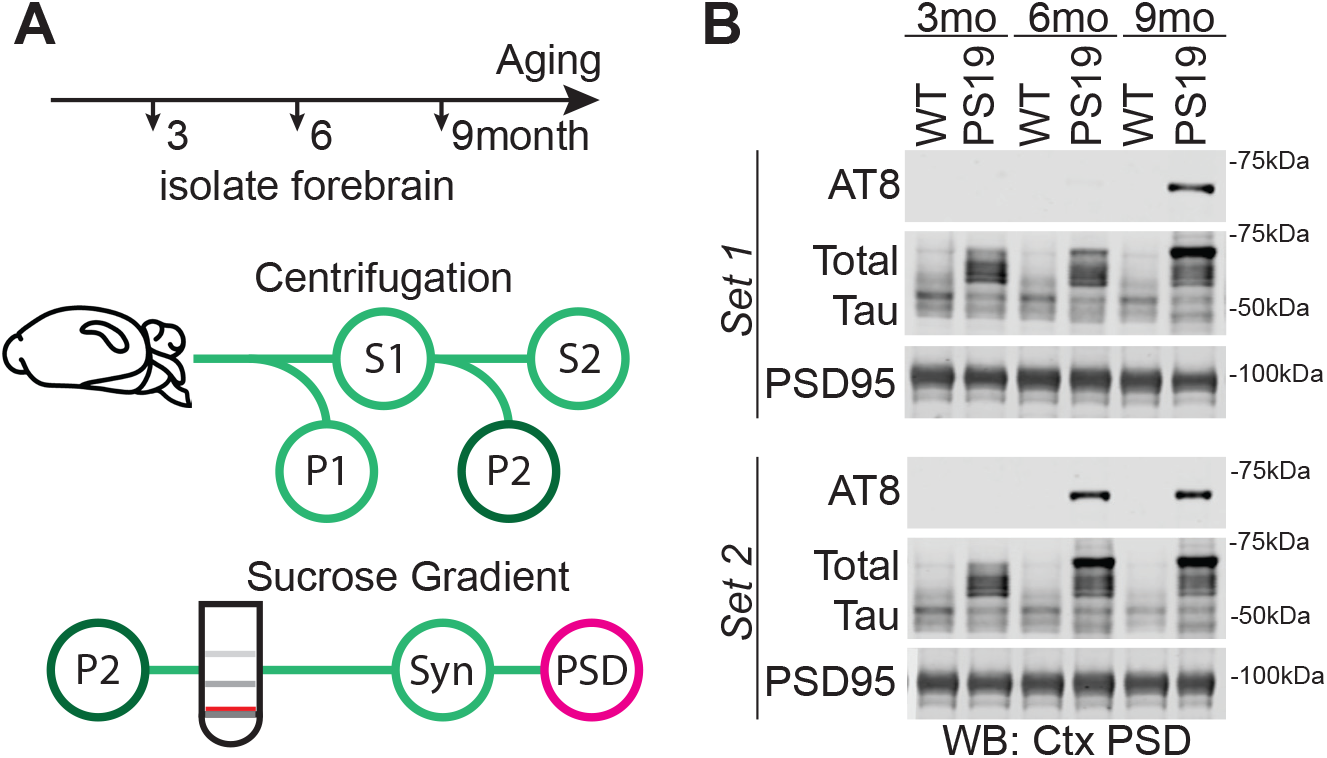
Tau in the cortex Post synaptic density (PSD). (A) Subcellular fractionation technique to isolate PSD. Red line indicates synaptosome fraction used to further isolate the PSD. (B) Western blot analysis showing phosphorylated Tau (AT8) accumulation with age in females, total Tau and PSD95, a protein enriched in the PSD.

Consistent with a prior report^23^, AT8-Tau was detected in forebrain PSD fractions in every PS19 mouse examined at 9months, whereas no PS19 mice were AT8-positive at 3months; example Western blots from females shown in Fig 2B. Interestingly, variable synaptic Tau burden was seen at 6 months in the cortex (Fig 2B) compare “set1” and “set2” in Fig. 2B. Approximately 50% of PS19 mice show AT8-Tau in forebrain PSD at 6months (Fig. 2B), and the remainder we conclude have not yet developed synaptic Tau burden. Comparable results were obtained from males (not shown). This observation suggested that 6months may be an age of particular vulnerability, where forebrain synaptic Tau burden is first beginning to appear, and where sleep disruption is already significant in both sexes (Fig. 1). Therefore, we centered our subsequent experiments on the 6month age in order to examine the possibility of a causal relationship between sleep disruption, Tau pathology, and cognitive decline.

### Sleep disruption is not causal in driving Tau burden in forebrain synapses

Because dark phase sleep disruption (hyperarousal) is clearly apparent at 6 months of age in both males and females, well in advance of the robust and widespread Tau pathology at 9months^23^ (Fig. 2), we hypothesized that sleep disruption may be a direct driver of Tau pathology in the forebrain synapse, as has been demonstrated in the brainstem LC neurons^25^. As a first step to test this hypothesis we examined whether variations in sleep behavior in PS19 mice at 6months are predicative of phospho-Tau in forebrain PSD fractions. Cohorts of 6 months old male and female PS19 mice underwent a 7day sleep recording, followed by sacrifice, isolation of the cortex PSD fraction, and Western blot analysis of several markers of Tau pathology. In order to provide a reliable standard for which to quantitatively compare Tau burden in our 6month old individuals, we pooled cortex PSD factions from 4x 11.5month (late-stage disease) PS19 mice of both sexes and included this late stage pooled PSD material in our Western blots. Tau burden in 6month old individuals, assessed through Western blot of a number of Tau epitopes including AT8, was then normalized to this late-stage 11month standard, and synaptic Tau burden was correlated with dark phase sleep measures for each mouse. Counter to our hypothesis, we found no statistical correlation (Pearson’s) between dark phase sleep amount, or bout length and synaptic AT8-Tau pathology in the cortex in females (Fig 3A-C) or males (Fig 3D-F). No other Tau epitopes were found to correlate with dark phase sleep measures (Supplementary Figure 1). These results show that sleep behavior immediately before sacrifice in 6-month-old PS19 mice is not predictive of synaptic Tau burden in the cortex.

**Figure 3.**
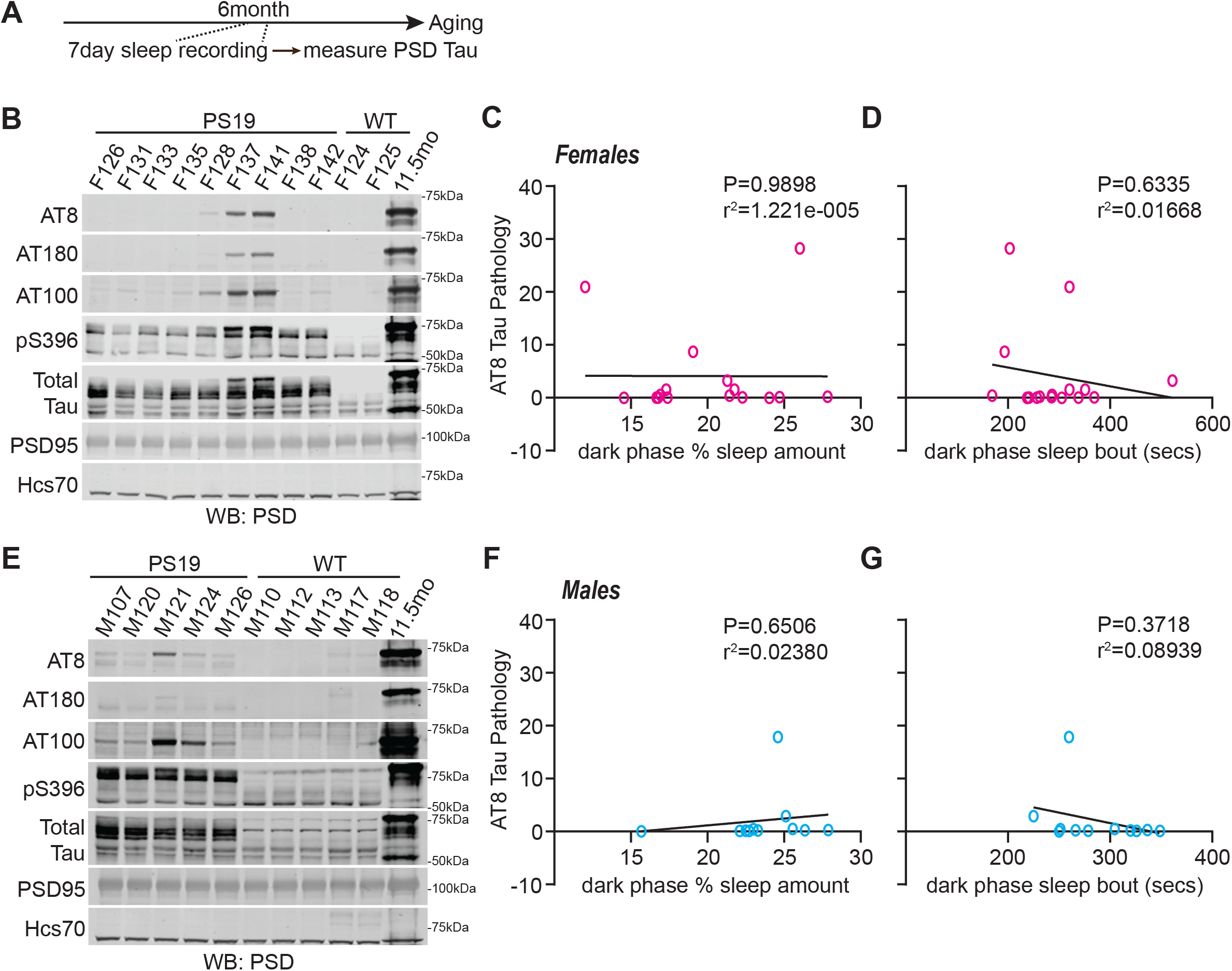
Decreased sleep amount in PS19 Tau tg mice is not predicative of AT8 Tau pathology in the cortex. (A) experimental design. (B) Western blot analysis of AT8, AT180, AT100, pS396, Total Tau, and PSD95 in 6-month (early phase) PS19 females. (C and D) Correlation of AT8 Tau pathology expression in the cortex of PS19 females with average dark phase hourly sleep (C) or sleep bout length in seconds (D). Sleep data was separated into 12hrs of dark and light phases. Dark phase sleep is represented here. N=16 PS19 females. (E) Western blot analysis of AT8, AT180, AT100, pS396, Total Tau, and PSD95 in 6-month PS19 males (F and G) Correlation of AT8 Tau pathology expression in the cortex of PS19 males with average dark phase hourly sleep (F) or sleep bout length in seconds (G). Sleep data was separated into 12hrs of dark and light phases. Dark phase sleep is represented here. N=11 All antibodies normalized to loading control. No significance (Pearson correlation).

Declines in sleep amount was previously shown to be correlated with Tau pathology in sleep promoting regions of the brainstem, but not in the cortex^24^, similar to our findings in Fig. 3. Experimentally induced sleep disruption has been demonstrated to be a direct driver of Tau pathology in the locus coeruleus (LC)^25^, but whether sleep disruption is a direct driver of Tau pathology at the forebrain synapse is unknown. To further examine the relationship between sleep disruption and forebrain synaptic Tau burden we tested whether experimentally induced acute 4hr total sleep deprivation (SD4) or 30days chronic sleep disruption (CSD) was causal in driving Tau pathology at the synapse at the vulnerable 6month age. Acute SD4 addresses the immediate response to sleep loss, while CSD investigates the accumulated consequences of sustained sleep loss/fragmentation. We, therefore, performed SD4 and CSD experiments on male and female PS19 mice and measured synaptic Tau burden in comparison to undisturbed controls, immediately following treatments at 6months of age. SD4 was achieved through gentle handling to keep mice awake for 4hrs starting from light onset (zeitgeber time, ZT0-4); control groups were left undisturbed and sacrificed at the same time of day (ZT4) (Fig. 4A). CSD is achieved by placing the home-cage on top of on an orbital shaker for 30days set to agitate at 110RPM for 10sec every 109sec (99sec rest) for 24hrs/day, providing a mild but regular stimulus demonstrated to cause sleep fragmentation^25, 35, 37^; control groups were housed in the same room and placed onto identical orbital shakers that were left off (Fig. 5A). Control or CSD treated mice were all sacrificed at ZT4. At the conclusion of SD4 or CSD, animals were sacrificed, and synaptic Tau burden was assessed using subcellular fractionation and Western blot. Compared to undisturbed controls, SD4 treatment had no effect on synapse Tau pathology in the cortex or hippocampus in PS19 mice of either sex (Fig. 4B-E). Surprisingly, the more sustained CSD treatment also had no measurable effect on cortex or hippocampus synaptic Tau burden in comparison to control treatments in either sex (Fig. 5B-E). Based on these findings we conclude that sleep disruption is not a direct driver of synaptic Tau burden in the forebrain, in contrast to what has been documented in the LC^25^.

**Figure 4.**
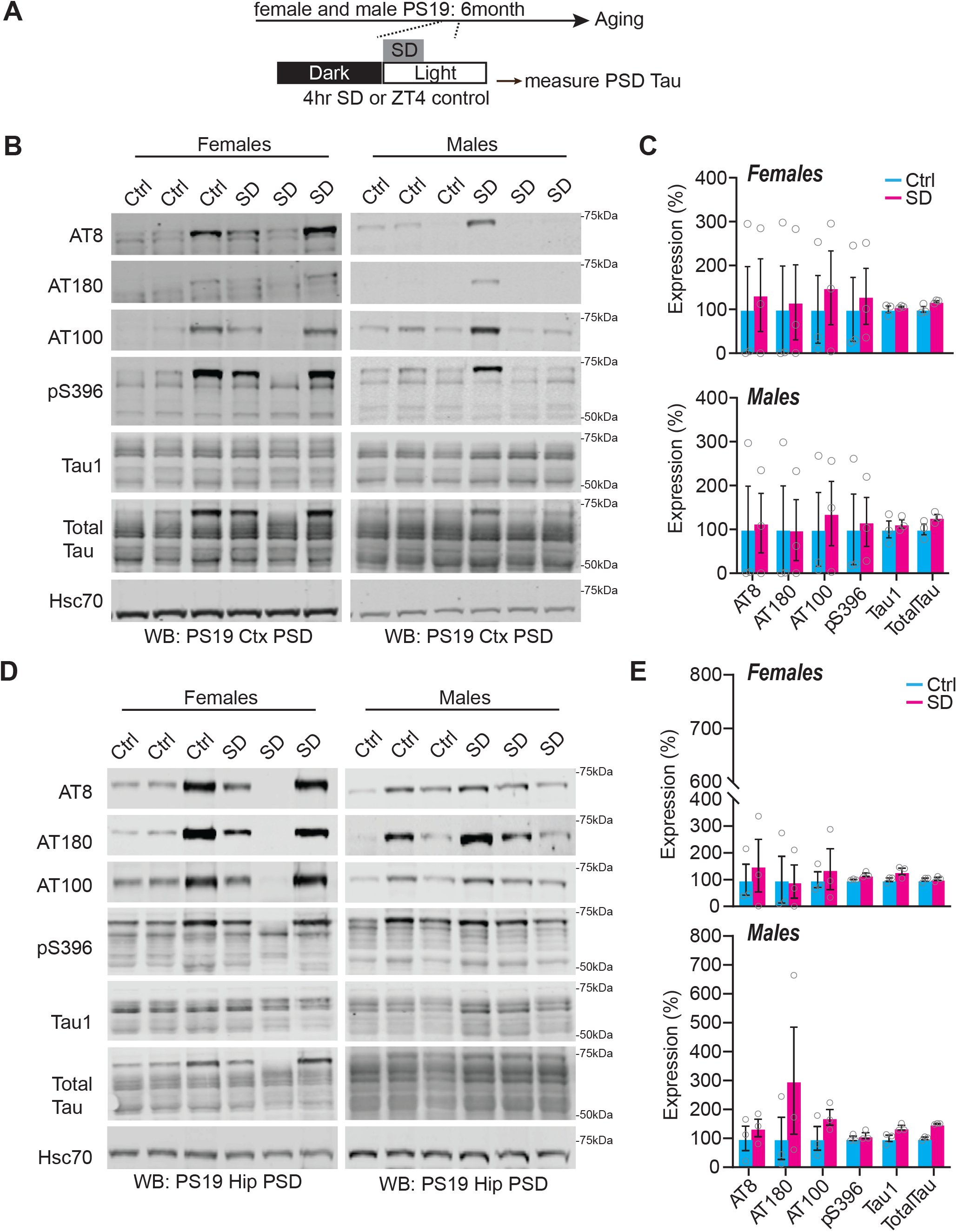
Synaptic Tau is not increased in PS19 mice after acute sleep deprivation in the cortex and hippocampus. (A) experimental design. (B) Western blot analysis of AT8, AT180, AT100, pS396, Tau1, and Total Tau cortical expression in 6-month (early pathology) PS19 females and males. (C) Quantification of cortical synaptic Tau proteins in PS19 females and males. N= 3 control; 3 CSD per sex. (D) Western blot analysis of AT8, AT180, AT100, pS396, Tau1, and Total Tau hippocampal expression in 6-month (early pathology) PS19 females and males. (E). Western blot analysis of AT8, AT180, AT100, pS396, Tau1, and Total Tau hippocampal expression in 6-month (early pathology) PS19 females and males. N= 3 control; 3 SD per sex. All antibodies normalized to loading control then normalized to control group. Unpaired two- tailed student’s t-test between control and treatment group. Error bars indicate mean ± SEM.

**Figure 5.**
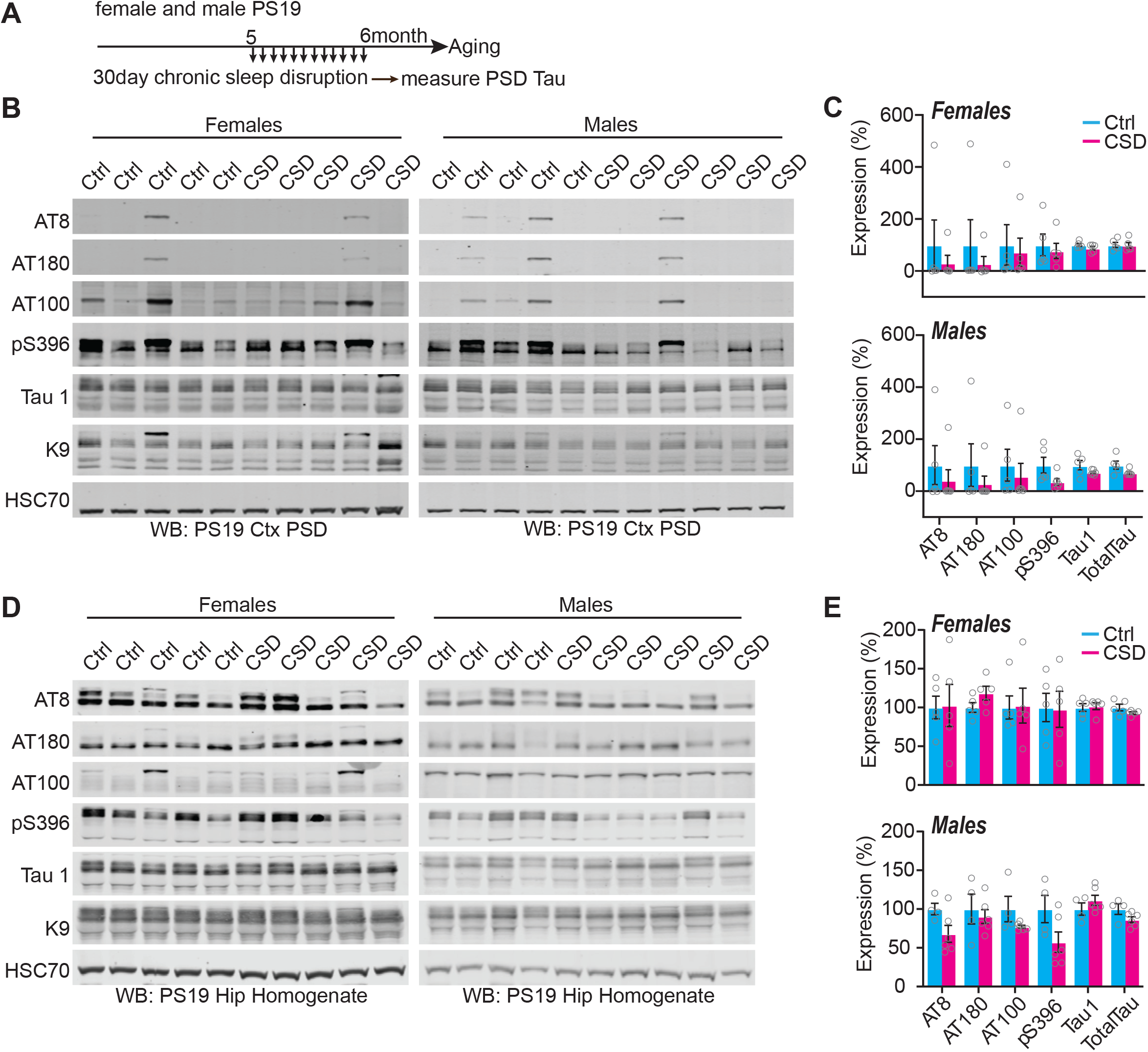
Synaptic Tau is not increased in PS19 mice after chronic sleep disruption in the cortex and hippocampus. (A) experimental design. (B) Western blot analysis of AT8, AT180, AT100, pS396, Tau1, and Total Tau in cortical PSD fractions of 6-month (early pathology) PS19 females and males. (C) Quantification of cortical synaptic Tau proteins in PS19 females and males. N= 5 control; 5 CSD females; N= 5 control; 6 CSD males. (D) Western blot analysis of AT8, AT180, AT100, pS396, Tau1, and Total Tau in hippocampus of 6-month PS19 females and males. (E) Quantification of hippocampal Tau proteins in PS19 females and males. N= 5 control; 5 CSD females; N= 4 control; 6 CSD males. All antibodies normalized to loading control then normalized to control group. Unpaired two-tailed student’s t-test between control and treatment group. Error bars indicate mean ± SEM.

### Chronic sleep disruption in PS19 drives female-specific adaptation in the hippocampus

Chronic sleep disruption has been shown to negatively impact AD pathogenesis and may leave synapses vulnerable to disease over time^4, 5^. While sleep disruption did not directly drive forebrain synaptic Tau pathology, we reasoned that the presence of synaptic Tau in PS19 mice may alter how synapses adapt to sleep loss, and thereby synergize with sleep loss to drive cognitive impairments. Therefore, we investigated whether cortex and hippocampus synaptic protein expression was altered in the male and female PS19 mice exposed to CSD treatment for 30days from 5-6months of age (as in Fig. 5A) in comparison to undisturbed controls. We focused on AMPA and NMDA-type glutamate receptors as these receptors are major mediators of synaptic strength and plasticity^38^. In addition, we examined the phosphorylation of AMPAR subunit GluA1 at two sites, S831 and S845, known to be involved in synaptic plasticity^38, 39^. Interestingly, in response to CSD treatment, female PS19 mice showed a striking increase in the expression of hippocampal AMPA and NMDA receptor subunits, GluA1 phosphorylation, as well as immediate early gene Arc and synaptic scaffold protein PSD95 in comparison to undisturbed controls (Fig. 6A-B). This response was completely absent in male PS19 mice (Fig. 6C-D). In contrast to our findings from hippocampus, no statistical differences were observed in the cortex of male or female PS19 mice exposed to CSD (Fig. 6). These results suggest that PS19 females, and not males, respond to chronic sleep disruption with a compensatory increase in synaptic protein expression in the hippocampus, but not the cortex. This result motivated us to investigate how CSD treatment affected hippocampal-dependent learning and memory in PS19 mice of both sexes.

**Figure 6.**
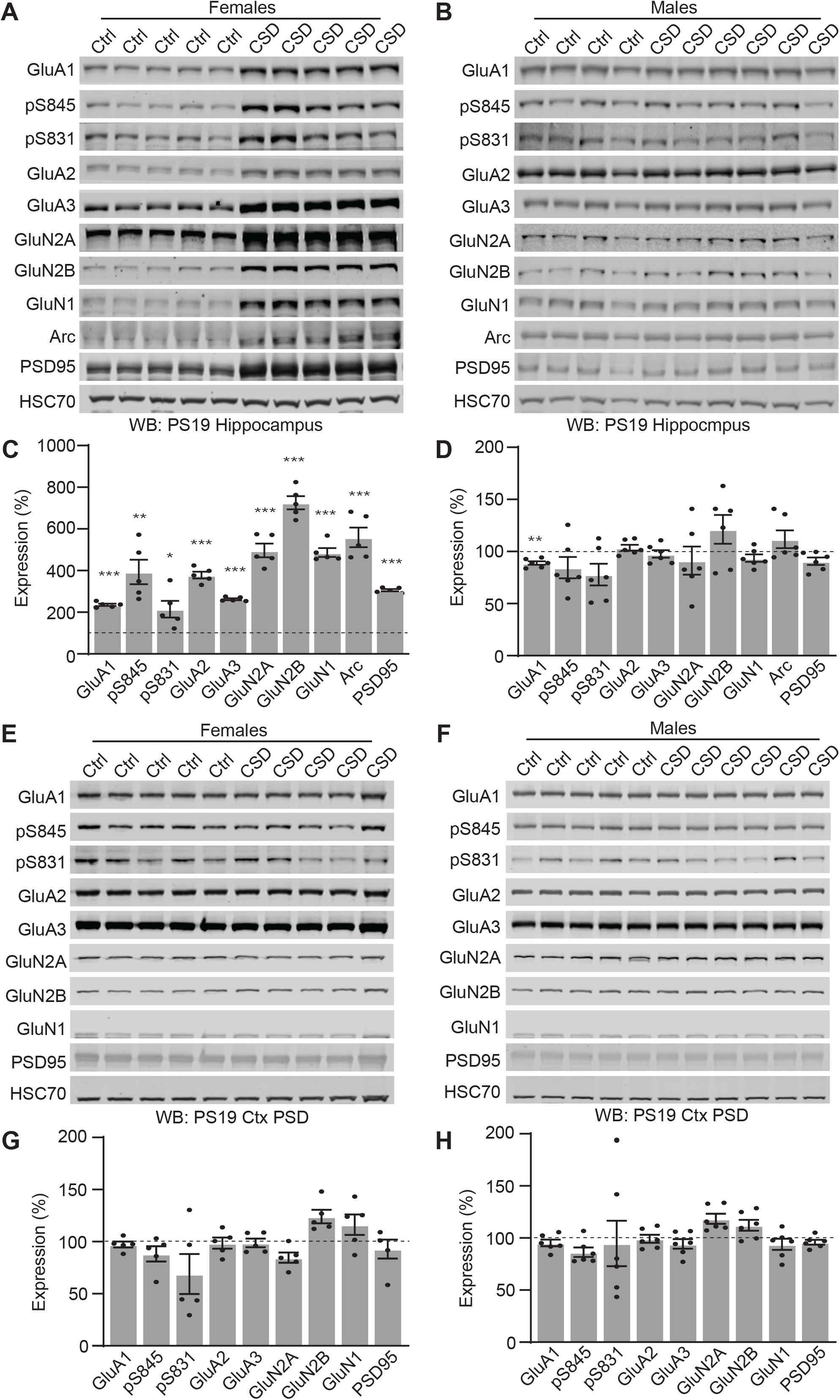
Chronic sleep disruption leads to changes in hippocampal synaptic protein expression in PS19 females but not males. (A, B) Western blot analysis of hippocampal synaptic receptor expression in 6-month PS19 females (A) and males (B). (C) Quantification of hippocampal protein expression in PS19 females. N= 5 control; 5 CSD. (D) Quantification of hippocampal protein expression in PS19 males. N= 4 control; 6 CSD. All antibodies normalized to loading control then normalized to control group. Unpaired two-tailed student’s t-test between control and treatment group for each genotype. *p<0.05, **p<0.01, ***p<0.001. Error bars indicate mean ± SEM. (E, F) Western blot analysis of cortical synaptic protein expression in 6-month PS19 females (E) and males (F). (G) Quantification of cortical synaptic protein expression in PS19 females. N= 5 control; 5 CSD. (H) Quantification of cortical synaptic protein expression in PS19 males. N= 4 control; 6 CSD. All antibodies normalized to loading control then normalized to control group. Unpaired two-tailed student’s t-test between control and treatment group for each genotype. Error bars indicate mean ± SEM.

### Chronic sleep disruption accelerates the decline of hippocampal dependent spatial learning and memory in male PS19 mice

Finally, we went on to investigate whether CSD, terminating at the 6-month vulnerable time point could accelerate the onset of cognitive decline. PS19 mice are known to display age-related deficits in learning and memory which can be accelerated via chronic sleep disruption treatments^25^. However, whether sleep disruption drives sex-specific effects on cognitive performance has not been investigated. We hypothesized that CSD during the vulnerable 6month age point would accelerate the onset of cognitive decline in this mouse model. Moreover, based on our molecular results showing differential responses to CSD in the hippocampus between males and females, we anticipated that there may be sex-specific vulnerabilities to the lasting effects of sleep loss in our PS19 mice. 5-month-old male and female WT and PS19 mice underwent CSD treatment for 30 days (as described above) followed by cognitive assessment via Morris Water Maze (MWM), behavior concluding at ∼7months (Fig. 7A). Note that this age is well in advance of cognitive impairment previously demonstrated in PS19 using MWM at 9months of age^23^. In the first phase of this testing, mice were placed into the water tank with the escape platform visible. Because PS19 mice are known to undergo age-related motor impairments^40^, we first confirmed that all mice tested were able to swim and locate the visible platform within 1min. All our mice were able to perform the task (not shown) and no mice were excluded from further testing. In the next phase of the experiment the platform was hidden, and mice underwent 4 days of training, 4 trials per day, to learn the location of the hidden platform based on spatial cues placed around the water maze. All our mice showed the expected reduction in escape latency (finding the hidden platform) with repeated days of training, with the exception of PS19 males exposed to CSD that showed significantly longer escape latencies than WT siblings, and no clear trend to reduce latency with multiple days of training, indicating that male CSD-PS19 mice show impairments in learning in this spatial task (Fig. 7B,D). In the final phase of the test, the hidden platform is removed, and the mice are returned to the water maze and their time spent searching in the former location of the platform is measured, a readout of spatial memory retention. Consistent with the above results all our mice spent significantly more time in the target quadrant of the maze than in the opposite quadrant indicating intact spatial memory, with the exception of male CSD-PS19 mice, indicating impaired memory retention in this group (Fig. 7C,E). Previous studies have shown that PS19 mice eventually develop impairments in the Morris water maze at 9months^23^, however, at the age tested here (6-7months), PS19 mice of both sexes in the control group performed comparably to WT littermates. Therefore, these results show that CSD treatment was able accelerate the onset of cognitive decline in male PS19 mice, whereas female PS19 and WT littermates were found to be resilient to these negative effects. We conclude that sleep disruption in concert with Tau pathology is a driver of cognitive decline, and that the sexes show differential vulnerability to these effects.

**Figure 7.**
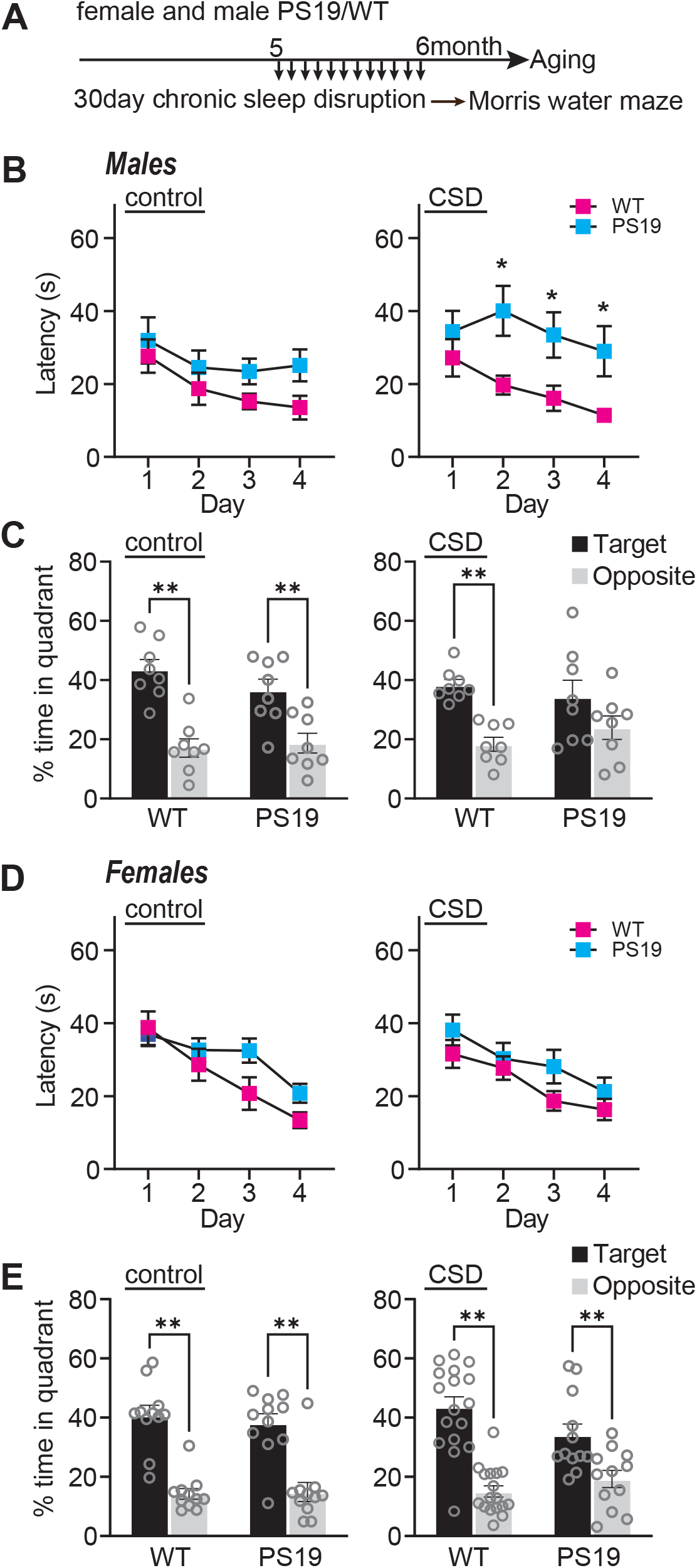
Chronic sleep disruption accelerates cognitive decline in males. (A) experimental design. (B) Males: Acquisition of spatial learning. Escape latencies during training in control and CSD treated mice. (C). Males: Spatial memory retention during 1min probe trial, time in target or opposite quadrant in control and CSD treated mice. Target indicates the quadrant where the platform had been located, versus the opposite quadrant. (D) Females: Acquisition of spatial learning. Escape latencies during training in control and CSD treated mice. (E). Females: Spatial memory retention during 1min probe trial, time in target or opposite quadrant in control and CSD treated mice. N=8-17 per group (sex, genotype, treatment). Data are means (± SEM). *p<0.05, **p<0.01.

## Discussion

Previous studies have shown that sleep disruption is present in human AD patients and animal models and that sleep disruption can contribute to Tau accumulation and spread^4, 5, 21, 25^. It is unknown whether the presence of synaptic Tau may affect restorative synapse remodeling during sleep or whether sleep disruption drives Tau pathology in the synapse. Synapses play a significant role in the restorative benefits of sleep^1, 7–10^. In the current study we examined the age-related progression of sleep phenotypes in relation to forebrain synaptic Tau burden in both male and female PS19 mice. We show that PS19 mice exhibit an early onset hyperarousal phenotype: reduced sleep time in the dark phase in comparison to WT littermates. Hyperarousal showed earlier onset in females, and in both sexes was progressive with age. At 3 months there is no AT8-positive Tau detected in the synapse fraction in any mice tested whereas at 9 months synaptic AT8 is detected in all PS19 mice tested. At 6 months approximately 50% of PS19 mice show synaptic AT8 indicating that this age may be a vulnerable timepoint where synapse-associated Tau pathology in the forebrain is emerging. We hypothesized that sleep disruption may be a direct driver of synaptic Tau pathology at this vulnerable 6month time point. However, counter to our hypothesis, we were not able to establish any correlation or causal relation between sleep and forebrain synaptic Tau burden. Strikingly, after CSD, PS19 females showed increased hippocampal synaptic protein expression not seen in males. Based on these data we hypothesized that CSD would drive acceleration of cognitive decline in hippocampal-dependent learning and memory in a sex-specific manner. Indeed, results of the Morris water maze show that chronic sleep disruption resulted in a clear acceleration of cognitive decline in male PS19 mice whereas females were found to be resilient. Our findings highlight two areas of interest for future studies. First, what mechanisms drive synaptic Tau pathology and does promotion of sleep mitigate consequences of synaptic Tau? Second, what mechanisms differentiate male and female responses to sleep disruption in AD, and what are the therapeutic implications of sex differences in the treatment of AD?

### Tau, Sleep deficits, and hyperarousal

Does Tau pathology drive sleep disruption, or does sleep disruption drive Tau pathology? A previous study used EEG to demonstrate that male PS19 mice show a progressive decline in sleep amount and bout lengths (sleep fragmentation) at 9-11months^24^, suggesting that sleep fragmentation may be a late-stage consequence of Tau pathology. In this prior study, sleep amount was found to negatively correlate with AT8-Tau pathology in sleep promoting regions of the brainstem, the sublaterodorsal area and parafacial zone, but not in the cortex, suggesting that sleep disruption may be caused by Tau pathology in these regions^24^. Our current data using non-invasive piezoelectric-based measures of home-cage sleep indicate that PS19 mice of both sexes exhibit sleep fragmentation at 9-11months (Fig. 1), consistent with the prior study^24^. In addition, we see alterations in sleep behavior occurring earlier than previously reported in the form of a selective dark phase hyperarousal, that was significant in females as young as 3months, and became apparent in males by 6months. Early stage hyperarousal may not have been noted in the prior EEG-based study because the dark phase was not analyzed separately, and females, with their earlier onset, were not included. Although hyperarousal was limited to the dark/active phase, it still resulted in a significant decrease in total daily sleep at 3months in females and 6months in males and was progressive thereafter. Because hyperarousal is apparent well in advance of robust Tau pathology, it may be driven by pre-aggregate Tau, or highly localized Tau pathology in discrete populations of wake/sleep promoting neurons (see below). As an early symptom, we hypothesized that hyperarousal may be causally linked with subsequent Tau pathology in the forebrain synapse, however we found no correlation between sleep and synaptic Tau burden at 6months. In future studies it will be important to test whether hyperarousal is a driver of Tau pathology in sub-cortical brain regions.

Wakefulness is controlled by discrete populations of wake-promoting neurons, the best studied of which are the orexinergic/hypocretin neurons of the lateral hypothalamus (LH), and the noradrenergic neurons of the locus coeruleus (LC)^41^. In nocturnal species including mice, these wake-promoting brain regions are more active in the dark phase. Interestingly, we find that sleep amount and bout lengths were not affected during the light phase in 3-9month old PS19 of either sex. We speculate therefore, that sleep behavior itself may not be impaired per se in the younger PS19 mice, but that wake promoting brain regions may be hyperactive during the dark phase, driving the hyperarousal phenotype. In accord with this idea, it was recently shown that sleep fragmentation occurring as a result of “natural aging” was caused by hyperexcitability of the orexin-LH neurons^42^. The LC is one of the first brain regions to exhibit Tau pathology in human AD patients, or even seemingly healthy younger adults, and also shows early stage pathology in PS19 mice^25–27^. Experimentally induced sleep disruption has been shown to drive Tau pathology in the LC, accelerating the loss of LC neurons^21, 25^. This suggests that LC may be one of the first brain regions affected in Tauopathies, and one of the most vulnerable brain regions to sleep loss. Moreover, LC neurons innervate vast territories of the forebrain, and the spread of pathological Tau variants across neuronal synapses has been proposed to be mediated by LC axons^43^. It is debated whether Tau drives neuronal hypo-or hyperexcitability, but an emerging trend is that Tau may drive hyperexcitability in early stages of disease before eventually driving hypoexcitation as neurons begin to succumb to pathology^44–46^. We speculate that in PS19 mice, a pre-aggregate, bioactive form of Tau may drive hyperexcitability in wake promoting orexin-LH or LC neurons, driving the dark phase hyperarousal phenotype we report. As hyperarousal progresses, loss of sleep and hyperactivation of LC neurons may initiate a feedforward cycle driving Tau pathology and neuron loss in the LC, leading to further changes in sleep behavior and promoting cognitive decline, consistent with prior reports^25, 47^. Further research in this area is certainly warranted.

### Sleep changes and association with forebrain synapses

Pathological Tau is known to spread between neurons via synaptic connections. Importantly, Tau release from neurons is a function of the sleep wake cycle and is exacerbated by sleep disruption and heightened neuronal activity^21^. In healthy neurons Tau is localized to axons but becomes mis-localized to the dendritic compartment and post-synapse during disease progression^16, 23^. We speculated that Tau mis-localization to forebrain synapses may also be driven by sleep disruption. Counter to our prediction, we did not see a correlation between sleep measures and synaptic Tau pathology in the cortex. Moreover, acute 4hr sleep deprivation or 30days of chronic sleep disruption in 6month old PS19 mice did not result in any change in forebrain synapse associated phospho-Tau compared to undisturbed controls. These findings suggest that sleep disruption may not directly cause Tau mistargeting to synapses. Consistent with these findings, and mentioned above, Holth et al. reported that sleep amount was negatively correlated with Tau pathology in sleep promoting regions of the brainstem, but not in cortex^24^. One possibility is that our sleep manipulations were insufficient and that more severe sleep disruption treatments (longer total deprivation, or more sustained chronic disruption) may drive an increase in Tau pathology at the synapse. Another possibility is that sleep disruption may indirectly promote Tau pathology in synapses, for example sleep disruption is known to exacerbate neuroinflammation^4, 5^. We suggest that while sleep disruption does not seem to be causally related to forebrain synapse Tau burden, that sleep disruption may synergize with synapse pathology to contribute to cognitive decline and neuron loss. Further research on the interaction between Tau pathology and sleep loss in the forebrain is required to address this.

### Sex-specific difference in onset of, and response to sleep disruption

We began our study by asking when sleep disruption and Tau pathology occurs in PS19 mice in an age and sex-dependent manner. This design is in accordance with recent recommendations that experiments should include both sexes^48^, which has not been well reflected in past AD animal research studies. We noted numerous important sex differences throughout our study. First, PS19 females showed an earlier onset of hyperarousal (3months), compared males (6months). In response to CSD treatment, females, but not males, show a hippocampal specific adaptation involving an upregulation of AMPA and NMDA-type glutamate receptors and other synaptic proteins. Finally, we show that CSD treatment was able to accelerate the onset of cognitive decline as measured using the Morris water maze in male PS19 mice, whereas females were found to be resilient. It is possible that female-specific upregulation of hippocampal synaptic proteins in response to CSD may form the basis of a type of “cognitive reserve” that was able to protect cognition of female PS19 mice, whereas males, lacking this putative adaptive mechanism, may be more vulnerable to the negative consequences of sleep loss. We note that this exciting possibility is highly speculative, and considerably more work will be needed to fully support this idea. Of note, men have been shown to sleep more than women, as women experience unique sleep disruptions during child-rearing. Men, however, experience more sleep architecture disruptions that worsen with age^49–51^ . It can be hypothesized that females develop more adaptations for sleep loss and thus contributes to the disparity seen in the exaggerated progressive decline in sleep in males^52^. The current limited pool of AD therapies does not account for sex differences; however, our data provides robust evidence for recognition of sex as a biological variable that deserves considerable attention.

### Conclusions

We show that sleep disruption, in the form of dark-phase hyperarousal, is an early symptom of disease progression in PS19 Tauopathy model mice. Chronic sleep disruption accelerated the onset of cognitive decline in PS19 males. Females show an earlier onset of hyperarousal, however, females showed resilience to the effects of chronic sleep disruption. We further conclude, that while sleep disruption is not a direct driver of Tau pathology in the forebrain synapses, sleep disruption likely interacts with synaptic Tau pathology to drive cognitive decline.

## List of abbreviations

AD: Alzheimer’s Disease
AMPA: 
ANOVA: analysis of variants
CSD: chronic sleep disruption
EEG: electroencephalography
EMG: electromyography
FTD: frontal temporal dementia
LC: locus coeruleus
LH: lateral hypothalamus
NFT: neurofibrillary tangle
NMDA: N-methyl D-aspartate
NREM: non rapid eye movement
PLSD: protected least significant difference
PSD: post synaptic density
REM: rapid eye movement
RPM: revolutions per minute
SD: sleep deprivation
SEM: standard error of the mean
UNC: University of North Carolina
WT: wild type
ZT: zeitgeber time

## Declarations

### Ethics approval and consent to participate

Animal procedures were all approved by Institutional Animal Care and Use Committee of the University of North Carolina (UNC) and performed according to guidelines set by the U.S. National Institutes of Health.

### Consent for publication

not applicable

### Availability of data and materials

The datasets used and/or analyzed during the current study are included in this manuscript and supplementary materials and are available from the corresponding author on reasonable request.

### Authors’ contributions

SCM and KKJ generated and genotyped all the mice for this study, performed sleep recording analysis, completed all biochemistry experiments and analysis, and participated in intellectual design of the project, analysis of the results and preparation of the manuscript and figures. KMH and SSM performed the behavior assays and completed the associated data analysis. TJC provided the founder P301S mice for the colony and assisted with study design and data interpretation. GHD designed and supervised the project, prepared the figures, and wrote the manuscript.

## Acknowledgements

The authors thank lab members Julia Lord and Micheal Ye for technical assistance.

---

**Supplementary Figure 1.**
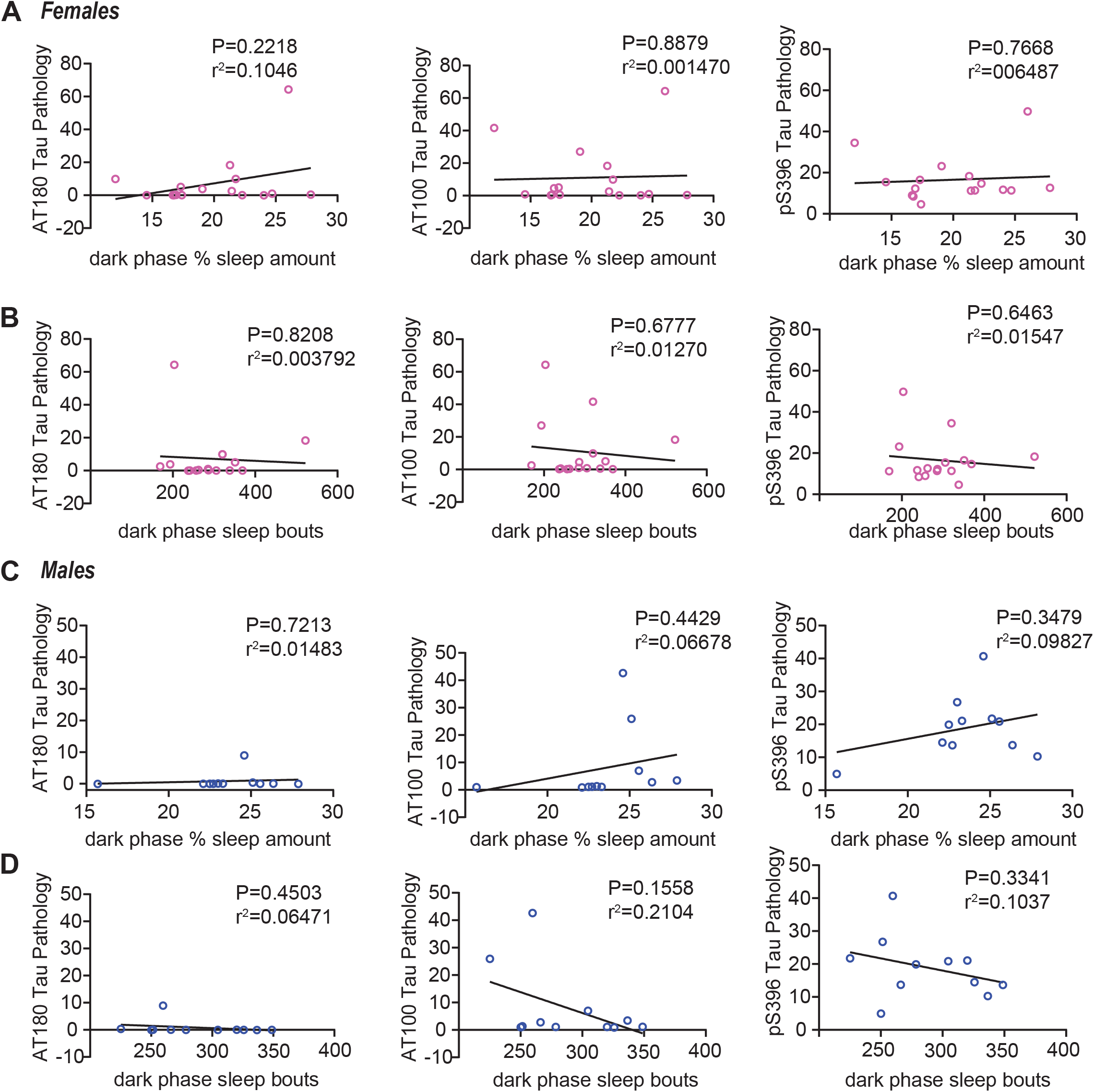
Decreased sleep amount in PS19 Tau tg mice is not predicative of AT8 Tau pathology in the cortex (continued). Western blots analysis of AT180, AT100, pS396 (see figure 3) correlated to dark phase sleep measures in 6-month (early phase) PS19 females and males. (A and B) Correlation analysis of AT180, AT100, pS396 Tau pathology expression in the cortex of PS19 females with average dark phase hourly sleep (A) or sleep bout length in seconds (B). Dark phase sleep is represented here. N=16 PS19 females. (C and D) Correlation analysis of AT180, AT100, pS396 Tau pathology expression in the cortex of PS19 males with average dark phase hourly sleep (C) or sleep bout length in seconds (D). Dark phase sleep is represented here. N=11 PS19 males. All antibodies normalized to loading control. No significance (Pearson correlation).

Supplementary Table 1. Summary sleep measures from Figure 1. Sleep amount: % of time spent sleeping in the light or dark phase. Average sleep bout length (s) in light and dark phase.

